# Experimental evolution of nascent multicellularity: Recognizing a Darwinian transition in individuality

**DOI:** 10.1101/2020.03.02.973792

**Authors:** Caroline J. Rose, Katrin Hammerschmidt, Paul B. Rainey

**Author notes:** Correspondence and requests for materials should be addressed to CJR.

## Abstract

Major evolutionary transitions in individuality, at any level of the biological hierarchy, occur when groups participate in Darwinian processes as units of selection in their own right. Identifying transitions in individuality can be problematic because apparent selection at one level of the biological hierarchy may be a by-product of selection occurring at another level. Here we discuss approaches to this “levels-of-selection” problem and apply them to a previously published experimental exploration of the evolutionary transition to multicellularity. In these experiments groups of the bacterium *Pseudomonas fluorescens* were required to reproduce via life cycles involving soma- and germline-like phases. The rate of transition between the two cell types was a focus of selection, and might be regarded as a property of groups, cells, or even genes. By examining the experimental data under several established philosophical frameworks, we argue that in the *Pseudomonas* experiments, bacterial groups acquired Darwinian properties sufficient to allow the evolution of traits adaptive at the group level.

## Introduction

Evolutionary transitions in individuality are central to the emergence of biological complexity. During the transition from single cells to multicellular organisms, groups of cells — on multiple occasions (Baldauf 2003) – have emerged as ‘Darwinian individuals’ capable of evolving by natural selection – as units of selection in their own right. The causes of these transitions are unclear. Nonetheless, philosophical and theoretical literature has provided insight into the fundamental requirements for evolutionary transitions to take place (Buss 1987; Maynard Smith and Szathmáry 1995; Michod 1999; Bonner 2000; Godfrey-Smith 2009; Mikhailov et al. 2009; Libby and Rainey 2013a; Shelton and Michod 2014; Black et al. 2020). Recently, the question has also received interest from the field of experimental evolution *(e.g.,* Ratcliff et al. 2012, 2013; Koschwanez et al. 2013; Hammerschmidt et al. 2014; Herron et al. 2019; Rose et al. 2020), which focuses on ecological circumstances that establish conditions promoting transitions to multicellularity. Here we attempt to bridge the gap between the fields of philosophy of biology and experimental biology by application of philosophical frameworks to experimental data.

## Levels of Selection

The view that the evolution of multicellularity necessitates a transition in individuality is based on a ‘multilevel’ perspective (Heisler and Damuth 1987; Damuth and Heisler 1988), which maintains that life is hierarchically organised, with ‘higher’ level units comprised of groups of particles from the level below. For example, eusocial colonies are made up of multicellular organisms, which in turn are integrated groups of cells. The very existence of a biological hierarchy is evidence of transitions from groups to Darwinian individuals. It follows that during the evolutionary history of the hierarchy, groups emerged as units that evolve by natural selection – that is, as entities in their own right that can evolve adaptations.

The essence of a transition in individuality lies in the capacity for higher-level reproduction, a prerequisite for Darwinian evolution. While group reproduction is necessary, it is not sufficient to ensure that natural selection operates at the level of the group. The archetypal ‘recipe’ for a trait to evolve by natural selection at a particular level of the biological hierarchy is that the trait under selection must be heritable, and it must covary with fitness (reproductive success), and must do so at that focal level (Lewontin 1970). However, covariance is a statistical concept and not a causal one. There *may* be a causal (selective) influence of a given trait on the fitness of a group if the trait covaries with group fitness, but covariance may also be misleading (Okasha 2006). Therefore, considering the possibility of natural selection independently at each level is inadequate because an apparent Darwinian process at one level can be a side effect of selection operating at another level – a ‘cross-level byproduct’ (Sober and Lewontin 1982; Okasha 2006; Shelton and Michod 2014). Okasha (2006, p.78-79) frames the ‘levels-of-selection problem’ as follows:

> *“When is a character^1^ -fitness covariance indicative of direct selection at the level in question, and when is it a by-product of direct selection at another hierarchical level?”*

Attempting to understand evolutionary origins of the biological hierarchy naturally leads to a multilevel view of evolution by natural selection, however alternative reductionist perspectives are also recognised. The supervenience argument, for example, maintains that genuine group-level selection is impossible because all properties of groups depend on those of their constituent particles. The traits of the underlying particles ultimately determine both the fitness and character of groups; the whole ‘supervenes’ on its parts (summarised in Okasha 2012). From such an argument it follows that there can never be a causal link between a group’s character and its fitness because *all* causality is at the particle level. The supervenience argument against group-level selection renders the levels-of-selection problem irrelevant because it makes cross-level by-products ubiquitous.

Perspectives that reduce all evolution to lower levels are not practical frameworks for investigating evolutionary transitions because of limited explanatory power. For example, the supervenience argument is unable to account for how the biological hierarchy itself came into existence and must logically reject the notion of a hierarchical structure in biological systems. Even the reductionist perspectives of Dawkins (1976, 1982) and others, which maintain that genic selection can account for all evolution, are not congruent with the supervenience argument because genes themselves are groups of lower level molecules whose evolution requires explanation. Okasha (2006) reasons that reductionist arguments only show that a trait-fitness covariance at the higher level must be a side effect of “*some lower level causal processes or other*, but not necessarily lower level *selection*”. Certainly everything in existence can be explained by its constituent particles, however reducibility to lower-level *selection* is what is important for understanding evolutionary origins. The underlying particle traits on which group fitness supervenes do not necessarily include particle *fitness,* so it does not follow that group fitness always has a lower-level *selective* explanation. Therefore, Okasha suggests that the concept of a cross-level by-product should be restricted to a trait-fitness covariance at one level that results from *selection* operating at another level.

In our view, if the level of *selection* is what matters, then characters or traits themselves should not be assumed to be an adaptation of any particular level prior to determining the level at which they have been selected. A trait may appear intuitively to be an adaptation of a particular level, but other cases are less clear. For example, a bistable switch between two cell phenotypes might be viewed as a property of cells, genes, or even a group of cells (manifesting as the proportion of the different cell phenotypes in the population). Even seemingly clear-cut cases may be less obvious when informed by the level of selection rather than by intuition. The trait ‘height’ is necessarily an attribute of some entity that exists at a particular level, *e.g.,* the height of a tree, or a termite mound. The height of a termite colony, however, may not play a role in selection between colonies, but could be a by-product of selection acting on individual termites within a colony for some trait that can only be empirically measured as the height of the mound. In other words, while the trait may intuitively be a property of one level, intuition cannot reliably determine if it is an *adaptation* of the level at which it is measured. The level at which the trait was selected should be determined empirically. The relevant factor in determining the level of selection is the level at which a given (undesignated) trait is causally associated with fitness.

It is possible to identify an evolutionary transition in individuality by determining the level(s) of selection. Groups evolve as higher-level Darwinian individuals when their evolutionary success (*i.e.*, fitness) can no longer be fully accounted for by the evolutionary success of their constituent particles. In other words, a primordial multicellular group constitutes a unit of selection once group functionality, rather than size, becomes the main determinant of fitness (Okasha 2006). Detecting the level of selection can be problematic because there exists a ‘grey area’ of mid-transition groups (Godfrey-Smith 2009). In such groups it is unclear whether certain properties are evolving due to selection at the group level, the cell level, or both. Importantly, traits should not be assigned to a particular level of selection based on assumed benefits to a given level. Furthermore, the emergence and evolution of Darwinian properties themselves (*i.e.*, reproduction and heritable variation) at the higher level requires a selective explanation. It is not sufficient to simply transpose the existence of Darwinian properties, such as variation, from the lower level to the higher (Griesemer 2000; Bourrat 2015); the possibility of selection acting upon this variation is dependent on the mode of group reproduction (Libby and Rainey 2013b; Rainey and de Monte 2014; De Monte and Rainey 2014; Hammerschmidt et al. 2014; Rose et al. 2020).

Hammerschmidt et al. (2014) reported the results of a selection experiment involving lineages of the bacterium *Pseudomonas fluorescens* that differed in the mode of group reproduction. Groups reproduced via a single-celled bottleneck in two selection regimes, however in one regime this bottleneck was an alternative cell type (a ‘germ’ cell). Crucially, while all groups exhibited the hallmarks of natural selection (reproduction, variation and heredity), improvements in group fitness became ‘decoupled’ from the fitness of constituent cells only for groups that reproduced via a germline.

This experiment illustrates some difficulties with a simplistic interpretation of the process of natural selection at multiple levels. Recognition of a transition in individuality calls for careful consideration of the possibility of cross-level byproducts. Various methods have been proposed for determining the hierarchical level(s) at which the association between a trait and fitness is causal, rather than merely statistical. Here we describe several approaches for tackling the levels-of-selection problem and apply them to the *P. fluorescens* populations. We demonstrate that a key trait evolved in the experiment due to selection at different levels depending on the environment.

## The Experiment

The Hammerschmidt et al. (2014) experiment tracked the evolution of groups of the bacterium *P. fluorescens*, which had previously been shown to evolve between matforming (soma-like) cells and dispersing (germline-like) states via negative frequency-dependent selection (Rainey and Rainey 2003; Libby and Rainey 2013b). When cultured in a static environment, rare cooperative “wrinkly spreader” (WS) mutants invade populations of planktonic “smooth” (SM) cells by forming a surface-colonizing soma-like mat that allows cells to access oxygen. Within the mat, SM “cheating” germline-like cells rise in frequency when rare due to advantages that stem from not producing costly cell-cell adhesive.

The frequency-dependent interactions between SM and WS cell types were utilized to implement a multicellular life cycle in which the WS mats were analogous to soma, and SM cells functioned as a germline. This facilitated differential group death and birth, which in turn allowed selection to operate between groups (Fig 1a). After ten life cycle generations, group fitness (relative number of group offspring produced) increased while cell fitness (total cell density) decreased; the fitness of the higher and lower levels became decoupled – a signature of a transition in individuality. Enhanced fitness of evolved groups was attributable to a property selected at the group-level, namely, the capacity to transition through the germ and soma phases of the life cycle (termed ‘transition rate’), and was not explained by improvements in individual cell proliferation – the fitness of the SM ‘propagules’ and the total carrying capacity were both reduced.

**Fig 1.**
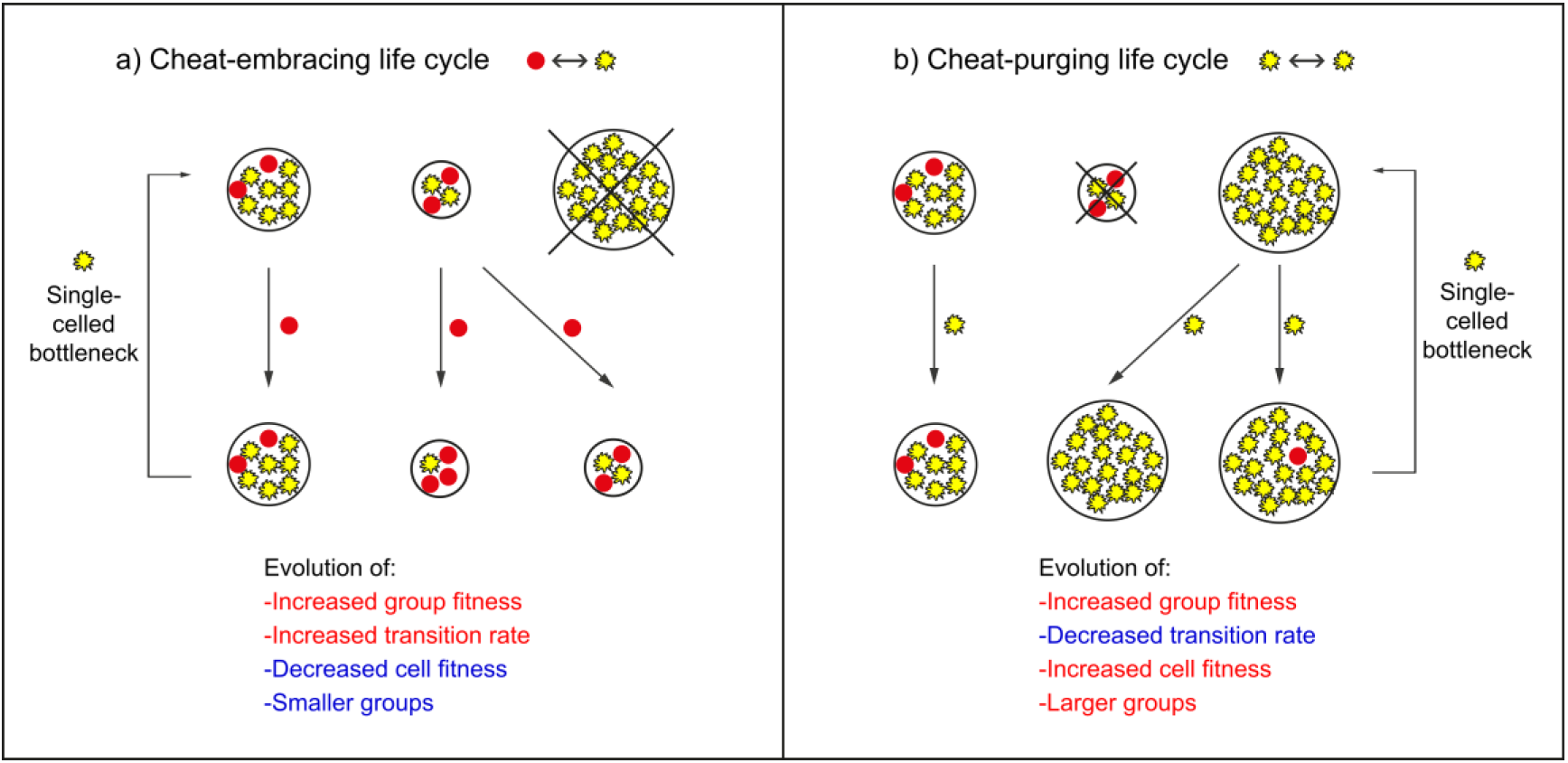
Simplified schematic contrasting the two experimental manipulations carried out by Hammerschmidt et al. (2014). a) In the “cheat-embracing” regime, groups of bacteria reproduced via a germline-like cell: a phenotypically different, non-cooperative cell type. b) In the “cheat-purging” regime, groups of bacteria reproduced via a single-celled bottleneck of the same cell type. Cell fitness is the total cell density, while group fitness is the relative ability of groups to beget offspring groups.

In parallel to this “cheat-embracing” selective regime, a “cheat-purging” experiment was conducted, in which groups reproduced via a single WS propagule cell (Fig 1b). While group fitness increased in both regimes, in the cheat-purging regime this was correlated with increases in all indicators of cell fitness. The following sections contrast the results of the cheat-embracing and cheat-purging ecologies in light of various perspectives arising from the levels-of-selection debate.

## Emergence

One approach to the levels-of-selection problem is to distinguish between an *emergent* trait – a property of a group that cannot be qualitatively or quantitatively reduced to the sum of its parts (Corning 2002), and *non-emergent* (or ‘aggregate’) traits. How such a distinction is made is not always straightforward because all properties of a group can ultimately be explained by the underlying particles (discussed above). Nevertheless, the ‘emergent character requirement’ (Vrba and Eldredge 1984; Vrba 1989) posits that genuine group-level selection, that is not reducible to selection at the particle level, can only operate on emergent traits. Okasha (2006) objects to this rule because an emergent trait may be a *result* of selection operating at the group-level, rather than a pre-condition of it. The capacity for group reproduction, which is necessary for group-level selection, is itself a derived, and arguably emergent, trait (Griesemer 2000; Rainey 2007; Rainey and Kerr 2010). It is also possible for an emergent trait to increase in frequency in the absence of group-level selection if it is associated with traits that increase particle fitness, *i.e.,* an emergent trait may inadvertently result from particle-level selection. It is therefore generally accepted that the emergent/aggregate distinction is not useful for solving the levels-of-selection problem (Damuth and Heisler 1988; Okasha 2006, 2012; Godfrey-Smith 2009).

Stephen Jay Gould argued that while a given trait need not be emergent to evolve at the level of the group, emergent *fitness* is what matters for group-level selection (Lloyd and Gould 1993; Gould 2002). It is true that groups must acquire the capacity for reproduction to allow the possibility of group-level selection and a measure of group fitness. However, this potential for group-level selection does not rule out cross-level by-products. Group fitness emerged (increased) in both the cheat-purging and cheatembracing selective regimes in Hammerschmidt et al. (2014), however in the cheatpurging regime this was likely due to a cross-level by-product because the emergent group fitness was correlated with increased cell fitness.

Heisler and Damuth (1987) suggest that emergence of the *relation* between a trait and group-level fitness is of fundamental importance for group-level selection. A traitfitness relation at a higher level is emergent if it cannot be accounted for by a traitfitness relation at a lower level. Okasha’s (2006) formulation of the levels-of-selection problem essentially recapitulates the emergent relation requirement. It is possible to identify an emergent relation when there is no positive association between particle and group fitness (Fig 2a,b).

**Fig 2.**
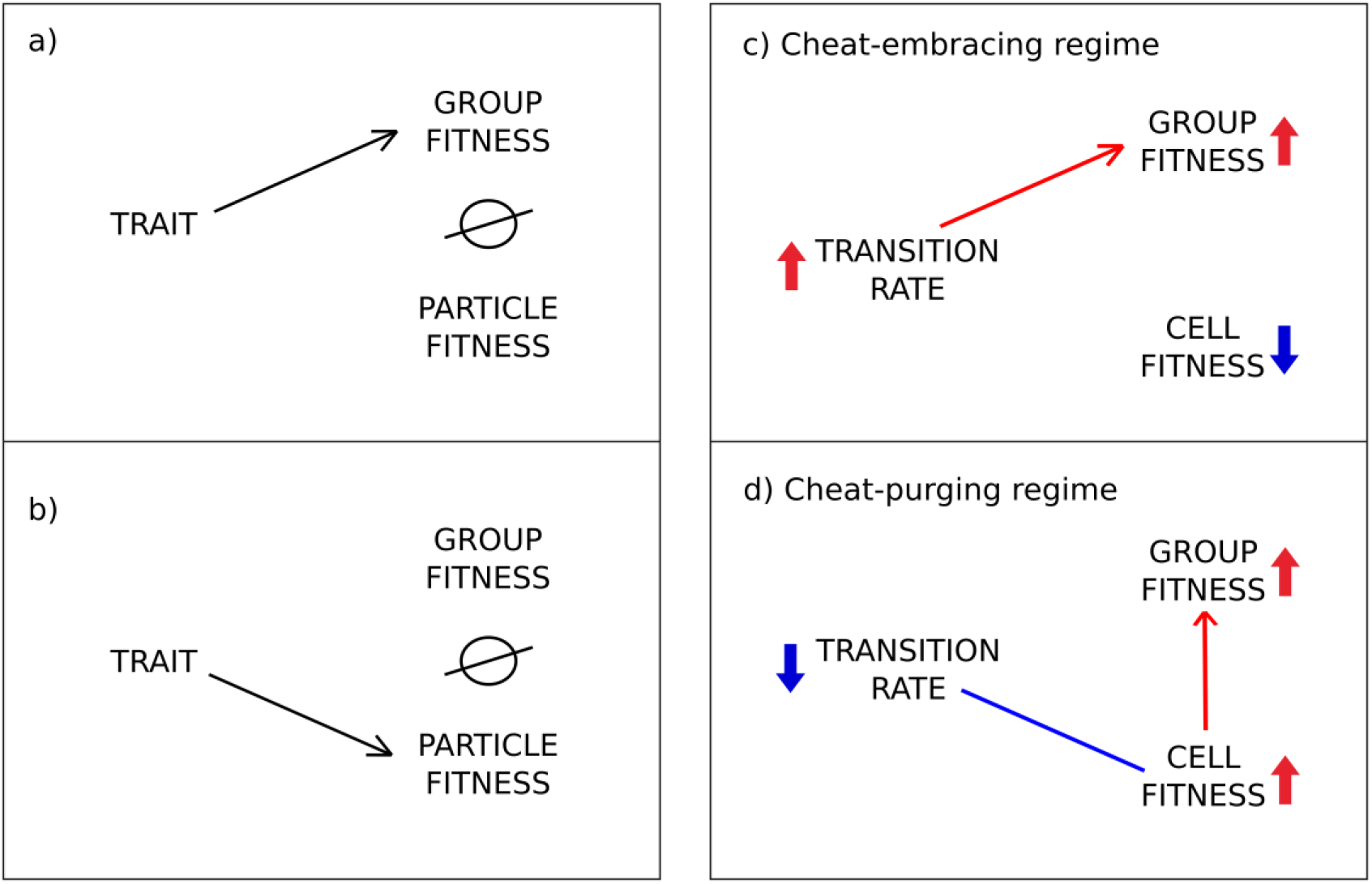
Emergent Relations. When there is no positive correlation between particle and group fitness, a trait-fitness relation at the group-level (a) is emergent; it cannot a by-product of a trait-fitness relation at the particle-level (b). c,d) Detection of Emergent Relations in the Hammerschmidt et al. (2014) experiment. In the cheatpurging regime (d) the transition rate trait correlates with cell fitness and not group fitness: transition rate decreased and group fitness increased as a result of cell-level selection. c) In the cheat-embracing regime, there is no correlation between cell fitness and group fitness: there is an emergent relation between transition rate and group fitness. Small red and blue arrows represent an increase or decrease in the parameters indicated relative to ancestral lines. Long lines represent positive (red) or negative (blue) correlations, and long arrows indicate a likely causal relationship.

A relationship exists between cell fitness and group fitness in the *P. fluorescens* groups that evolved under the cheat-purging regime (Hammerschmidt et al. 2014). This (probably causal) link presents no difficulty in determining the level of selection, because the trait in question (transition rate) correlates only with parameters of cell fitness, not group fitness (Fig 2d). It is clear therefore, that the transition rate significantly decreased because of selection acting at the cell level, and the increased group fitness can be attributed to the correlation between group and cell fitness parameters – group fitness supervenes on particle fitness. This is in contrast to the cheat-embracing regime, in which no correlation exists between cell and group fitness in the evolved populations thus the positive relationship between the increased transition rate and group fitness is causal – an emergent relation (Fig 2c).

## Contextual Analysis

Heisler and Damuth (1987) devised a method for detecting emergent relations by analysing trait-fitness covariance from phenotypic data. Contextual Analysis is a generalisation of the selection analysis developed by Lande and Arnold (1983). This approach treats the levels-of-selection problem as a special case of selection on correlated traits, where the dependence of fitness on correlated traits is captured with a standard linear regression model. The covariance of group fitness *(ϒ)* with a (so-called) ‘group’ trait (*Z*) could be non-zero *(i.e., Cov(ϒ,Z) ≠ 0*) in the absence of a causal link between *Z* and *ϒ* if *Z* correlates with another trait that *does* affect fitness. There is a cross-level by-product if *ϒ* is causally determined by the average fitness of its constituent particles, *W*. The trait-group fitness covariance can be partitioned into two components: direct group-level selection on a trait, *β_1_Var(Z)*, and the by-product of particle-level selection, *β*_2_Cov(W,Z):

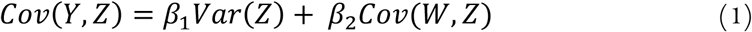

where *β*_1_ is a measure of the direct effect of *Z* on *ϒ*, controlling for *W*, and *β*_2_ measures the effect of *W* on *ϒ*, controlling for *Z*.

No relationship exists between transition rate and cell fitness in the cheat-embracing *P. fluorescens* populations (Hammerschmidt et al. 2014). Contextual analysis is therefore not necessary to partition the change in transition rate due to cell- and group-level selection, because *Cov(W,Z) = 0.* The strength of group-level selection for increased transition rate is equal to the covariance between group fitness and transition rate:

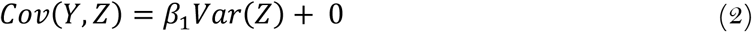

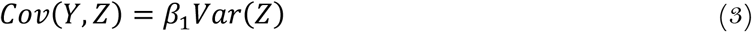

As no relationship exists between transition rate and group fitness in the evolved populations from the cheat-purging regime (Hammerschmidt et al. 2014), there is no need to discriminate between levels of selection – decreased transition rate was selected entirely at the lower level:

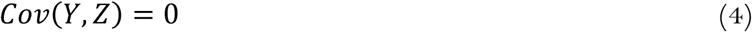

These results corroborate the idea that a trait should not assumed to be an adaptation of a particular hierarchical level. The transition rate between the two cell types might be regarded as a property of groups, cells, or even genes, often depending on the inclinations of the investigator. Our view is that transition rate should not be associated with one level because the direction of its evolution can change due to selection operating at multiple levels in differing ecological contexts (Rose et al. 2020). This trait evolved to be higher in the cheat-embracing selective regime as a result of its influence on group fitness, and lower in the cheat-purging regime because of selection at the cell-level. Does it then follow that transition rate is a group trait in one ecological context and a cell trait in another? To reiterate, the level of *selection* is the focus during an evolutionary transition in individuality. An *a priori* designation of transition rate as either a ‘group trait’ or a ‘cell trait’ may have masked the true level of selection, or worse, led to false conclusions.

## Additivity

Lloyd (1989) contributed the ‘additivity criterion’ to the levels-of-selection debate – a notion that was first put forward by Wimsatt (1980). The additivity criterion posits that true group-level selection can only occur when the variance in group fitness does not relate additively (linearly) to particle composition. When there is a linear relationship between group fitness and particle composition, then differences in group fitness can be fully explained by selection at the particle level. A shift in the level of selection occurs when there is a departure from additivity, *i.e.*, there is a non-linear relationship between group fitness and particle composition. The additivity criterion for group-level selection is similar to the criterion of emergent relations: a ‘new’ nonlinear relation between a trait and fitness emerges at the level of the group. However the additional specification that this relation be non-linear adds a crucial layer of information that the emergent relation requirement alone lacks.

The cheat-embracing selective regime in the *P. fluorescens* populations clearly results in a non-linear relationship between group fitness and particle composition. SM ‘germline’ cell types are required at a high enough frequency to be consistently detected during the regime, however a WS mat must also be present to avoid extinction. Therefore, group fitness will be highest at intermediate frequencies of SM cells, and zero if the proportion of SM cells is either 0% or 100%. This non-additive relationship between cell composition and group fitness is not pronounced in the experimental data (published in Hammerschmidt et al. (2014) but re-analysed here) because the wildtype transition rate is so low that the hypothetical ‘high SM/low group fitness’ groups do not exist in the ancestral populations, and were not selected during the experiment due to their low group fitness (Fig 3a). Nevertheless, a comparison of the fit of linear and quadratic models to the data confirms the non-additive relationship between the proportion of SM cells within a group and fitness. Akaike’s Information Criterion (Akaike 1973), which compares the quality of statistical models by giving a relative estimate of the information lost by overfitting a given model, indicated that the quadratic model was 99.99% and 99.88% more likely to explain the data than the linear model in the ancestral and evolved groups respectively (Supplementary Table). This non-additive, and therefore emergent, relationship between the proportion of SM cells (determined by the transition rate trait) and group fitness is indicative of selection operating at the level of the group.

**Fig 3.**
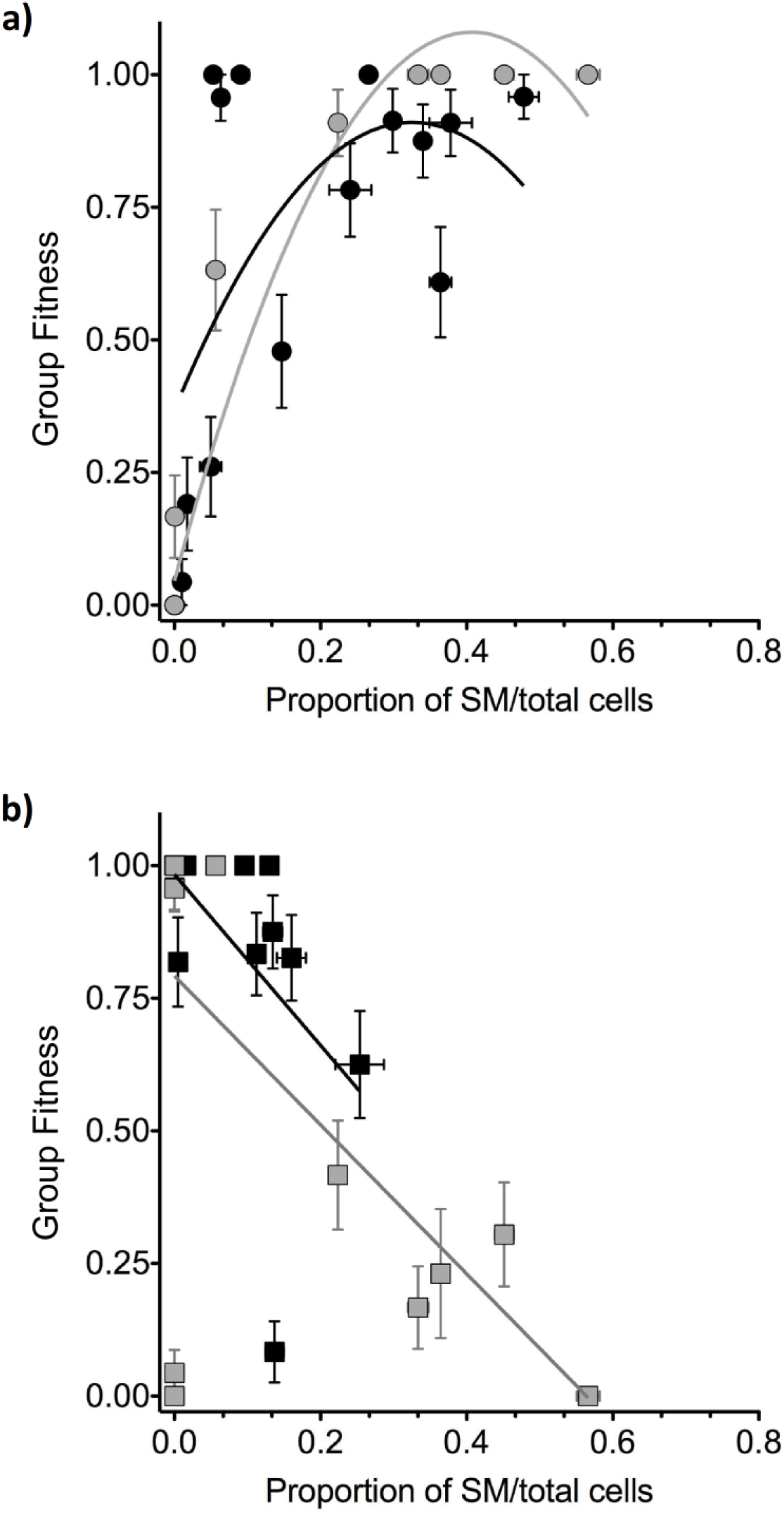
Relationship between particle composition and group fitness in the cheat-embracing (a) and cheat-purging (b) regimes. Ancestral = grey, evolved = black. Error bars are SEM, based on N=3.

In contrast, there is an additive relationship between the composition of groups (as determined by transition rate) and their fitnesses in the cheat-purging selective regime (Fig 3b). Compared to the quadratic model, the linear model has a 70.54% and 69.94% probability of correctly fitting the ancestral and evolved data respectively. The two ancestral lines that had both low group fitness and low proportion of SM cells did not exhibit visible mats because of the invasion of a third cell type, the “fuzzy spreader” (described in Ferguson et al. 2013). These groups were therefore extinguished after one generation and the proportion of SM alone became a reliable predictor of group fitness. In contrast to the non-linear relationship between the fitness of a group and its composition in the cheat-embracing regime, the additive dependence of a group’s fitness on its composition reveals lower-level causality in the cheat-purging regime. Non-additivity in the cheat-embracing life cycle led to a transition in individuality that was not possible for groups in the cheat-purging regime, whose fitnesses depended entirely on particle composition.

## Darwinian Populations

The essentialist search for ‘necessary and sufficient’ conditions to categorise evolutionary phenomena such as evolutionary transitions is often futile because of the dynamic nature of the biological world. Peter Godfrey-Smith (2009) (hereafter PGS) provides a highly intuitive method for visualising possible cases of Darwinian individuality as a gradient. This conceptualisation eliminates the need to quibble over whether any given biological example meets a certain threshold for individuality. PGS defines a Darwinian individual as a member of a Darwinian population: a collection of entities that undergo evolution by natural selection. A transition in individuality therefore involves the appearance a new kind of Darwinian population, which can exist in a gradient ranging from marginal, to minimal, to paradigm.

PGS emphasises the significant role of group reproduction in generating new levels of Darwinian individuals. The particular challenge of relevance to major evolutionary transitions is to identify which are cases of reproduction of groups, and which are simply cases of growth of groups resulting from multiplication and structural organization of their components. This view is related to the levels-of-selection problem, however, in framing the problem this way PGS recognizes that there are more than these two distinct alternatives and treats the possibility of group reproduction itself as a gradient of paradigm to marginal forms. During a transition in individuality, a marginal Darwinian population may initially be composed of group entities that have minimal capacity to leave offspring that resemble parental types, but are nevertheless able to evolve towards a more paradigm population.

PGS introduces a number of spatial tools that enable visualisation of the spectrum of possible Darwinian populations in a 3D ‘Darwinian Space’ representing the relationships between evolutionarily important parameters. The possibilities of different kinds of group reproduction can be visualised on a spatial illustration displaying three parameters that characterise paradigm examples of group reproduction (Fig 4). ‘*B*’ stands for ‘bottleneck’, and represents the extent of narrowing of a group during each generation – distinguishing reproduction from growth. The ‘germline’ parameter ‘*G*’ conveys the degree of reproductive specialisation of parts (a discussion on the importance of bottlenecks and germlines during multicellular reproduction is beyond the scope of this paper, and can be found elsewhere (Godfrey-Smith 2009; Rose 2020). The *P. fluorescens* groups in the experiments described in Hammerschmidt et al. (2014) have the highest possible value for *B,* a single-celled bottleneck, which is hypothesised to be a crucial factor for the evolution of individuality (Grosberg & Strathmann, 1998; Michod & Roze, 1999; Griesemer, 2000; Wolpert & Szathmáry, 2002). However, only groups subjected to the cheat-embracing selective regime have high values of *G,* while those in the cheat-purging regime have no reproductive specialisation: *G*=0.

**Fig 4.**
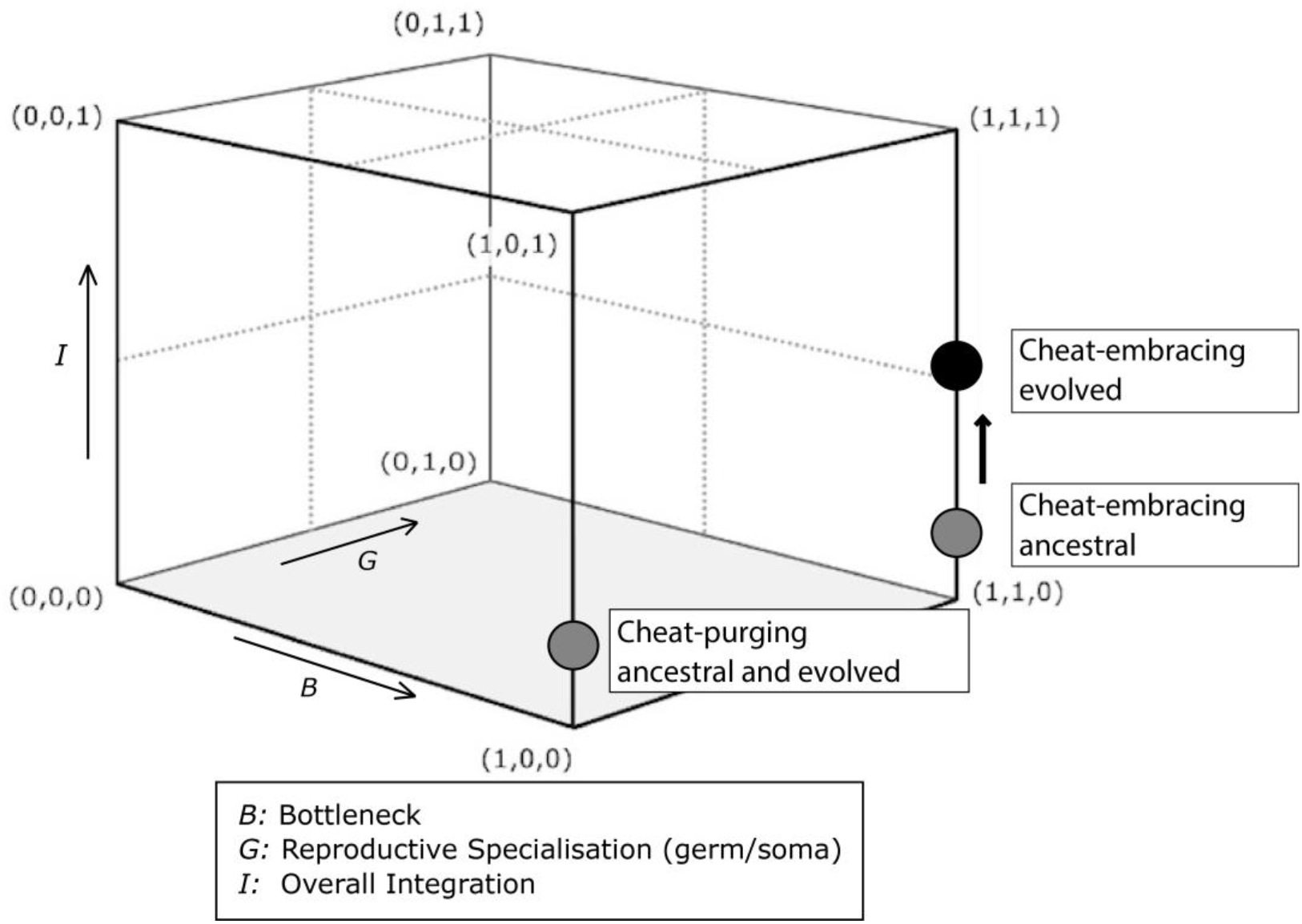
The spectrum of group reproduction in *P.fluorescens* groups. Groups that evolved in the cheat-embracing regime are closer to ‘paradigm’ (1,1,1) forms of reproduction than cheat-purging groups. Adapted from Godfrey-Smith (2009).

The ‘integration’ of the group is summarised in an overall sense by the *‘I’* parameter, which encompasses features such as the extent of non-reproductive division of labour, the mutual dependence (loss of autonomy) of parts, and the maintenance of boundaries surrounding groups. Both the cheat-embracing and cheating-purging groups have fairly low values of *I*. The cooperation required to form WS mats is a kind of integration in the form of mutual dependence of parts – though the parts that are dependent upon each other (WS cells) are identical. However, the selective advantage generated by the mat is a within-group advantage (access to oxygen denied to those cells outside of the mat) and therefore the mat does not contribute to the overall integration of a group. On the other hand, a mat is required in order to avoid extinction during both selective regimes, so in this sense a between-group selective advantage resulted from this basic level of group integration (*I*).

Subsequent to between-group selection, some *P. fluorescens* groups may have reached a higher level of integration (*I*) due to the evolution of developmental control. Hammerschmidt et al. (2014) describe the evolution of a genetic switch in the fittest of the cheat-embracing lines, line-17. A mutation in the mismatch repair locus *mutS* facilitated fast and reliable transitioning between the WS ‘soma’ and SM ‘germ’ phases of the life cycle via expansion and contraction of a tract of guanine residues in the diguanylate cyclase-encoding *WspR* – one of 39 genes known to underpin the WS phenotype (Spiers et al. 2002; Goymer et al. 2006; Lind et al. 2019). This genetic switch emerged within just ten group generations and upon extended selection may even come under developmental regulation. Visualising the *P. fluorescens* populations in this way makes it clear that while neither experimental regime resulted in the evolution of paradigm Darwinian populations, those groups that reproduced via a germline moved closer in ‘Darwinian Space’ towards individuality.

## Conclusion

The results from the experiments presented in Hammerschmidt et al. (2014) and scrutinized here in further detail demonstrate that a major evolutionary transition to multicellularity is possible when groups of cells reproduce via a two-phase life cycle involving a germline-like cell. Conflict resolution evolved in groups that reproduced without a germline (in the form of reduced transition rate to SM ‘germline-like’ cell types), but individuality did not manifest at the group level. In contrast, during the course of the evolution experiment, a causal relation between the transition rate trait and group fitness emerged in groups that reproduced via a germline-like cell. This emergent relation, in addition to a non-additive relation between the composition of the groups and group fitness, demonstrates that the levels-of-selection problem was resolved, shifting selection to the higher level and revealing new biological individuals that were closer to paradigm status in ‘Darwinian Space’. Importantly, while the transition rate trait evolved to be lower in one experimental regime due to selection acting at the cell-level, the same trait increased due to group-level selection in the other experimental setup, challenging the common practice of assigning traits to levels prior to rigorous levels-of-selection analyses. Primitive groups reproducing via a germline might eventually accumulate higher-level adaptations that lead to such complex integration of their comprising cells that they can no longer exist independently, surviving and replicating only as components of the group – the ‘organism’.

## Declarations

### Availability of data and material

Request for materials should be addressed to the corresponding author.

### Code Availability

Not applicable

## Supplementary Table

Comparison of the fit of linear and quadratic models of the relationship between the proportion of SM cells and group fitness in the cheat-embracing (CE) and cheatpurging (CP) regimes using the Akaike’s Information Criterion.

Prob correct: Probability that this model is the most likely to have generated the data.

| Parameters | Difference in AICc | N | Prob Correct | df | $R^2$ |
| --- | --- | --- | --- | --- | --- |
| **CE ancestral** | 151.5 | 360 |  |  |  |
| Linear model |  |  | <0.01% | 358 | 0.712 |
| Quadratic model |  |  | >99.99% | 357 | 0.812 |
| **CE evolved** | 13.41 | 336 |  |  |  |
| Linear model |  |  | 0.12% | 334 | 0.128 |
| Quadratic model |  |  | 99.88% | 333 | 0.167 |
| **CP ancestral** | -1.747 | 360 |  |  |  |
| Linear model |  |  | 70.54% | 358 | 0.291 |
| Quadratic model |  |  | 29.46% | 357 | 0.292 |
| **CP evolved** | -1.689 | 360 |  |  |  |
| Linear model |  |  | 69.94% | 358 | 0.141 |
| Quadratic model |  |  | 30.06% | 357 | 0.142 |

## Footnotes

1 A semantic note: The term ‘trait’ is used throughout the paper, however ‘character’ is on occasion used interchangeably with ‘trait’ when adopting the language of key reference material. ‘Group’ and ‘collective’ are also used interchangeably in this manner, as is the term ‘particle’, which is used by some references to refer to the units that comprise groups.

## References

Akaike H (1973) Information theory and an extension of the maximum likelihood principle. In: Petrov BN, Csaki F (eds) 2nd International Symposium on Information Theory. Akademiai Kiado, Budapest, pp 267–281

Baldauf SL (2003) The deep roots of eukaryotes. Science 300:1703–1706. https://doi.org/10.1126/science.1085544

Black AJ, Bourrat P, Rainey PB (2020) Ecological scaffolding and the evolution of individuality. Nat Ecol Evol 4:426–436. https://doi.org/10.1038/s41559-019-1086-9

Bonner JT (2000) First Signals: The Evolution of Multicellular Development. Princeton University Press, Princeton

Bourrat P (2015) Levels of Selection Are Artefacts of Different Fitness Temporal Measures. Ratio 28:40–50. https://doi.org/10.1111/rati.12053

Buss LW (1987) The Evolution of Individuality. Princeton University Press, Princeton

Corning PA (2002) The re-emergence of “emergence”: A venerable concept in search of a theory. Complexity 7:18–30. https://doi.org/10.1002/cplx.10043

Damuth J, Heisler IL (1988) Alternative formulations of multilevel selection. Biol Philos 3:407–430. https://doi.org/10.1007/bf00647962

Dawkins R (1976) The Selfish Gene. Oxford University Press, Oxford

Dawkins R (1982) The Extended Phenotype: The Gene as the Unit of Selection. Oxford University Press, Oxford

De Monte S, Rainey PB (2014) Nascent multicellular life and the emergence of individuality. J Biosci. https://doi.org/10.1007/s12038-014-9420-5

Ferguson GC, Bertels F, Rainey PB (2013) Adaptive divergence in experimental populations of *Pseudomonas fluorescens*. V. Insight into the niche specialist “Fuzzy Spreader” compels revision of the model *Pseudomonas* radiation. Genetics 195:1319–1335. https://doi.org/10.1534/genetics.113.154948

Godfrey-Smith P (2009) Darwinian Populations and Natural Selection. Oxford University Press, Oxford

Gould SJ (2002) The Structure of Evolutionary Theory. Belknap Press, Cambridge, MA

Goymer P, Kahn SG, Malone JG, et al (2006) Adaptive divergence in experimental populations of *Pseudomonas fluorescens*. II. Role of the GGDEF regulator WspR in evolution and development of the wrinkly spreader phenotype. Genetics 173:515–526. https://doi.org/10.1534/genetics.106.055863

Griesemer J (2000) The units of evolutionary transition. Selection 1:67–80. https://doi.org/10.1556/Select.1.2000.1-3.7

Hammerschmidt K, Rose CJ, Kerr B, Rainey PB (2014) Life cycles, fitness decoupling and the evolution of multicellularity. Nature 515:75–79. https://doi.org/10.1038/nature13884

Heisler IL, Damuth J (1987) A method for analyzing selection in hierarchically structured populations. Am Nat 130:582–602

Herron MD, Borin JM, Boswell JC, et al (2019) De novo origins of multicellularity in response to predation. Sci Rep 9:1–9. https://doi.org/10.1038/s41598-019-39558-8

Koschwanez JH, Foster KR, Murray AW (2013) Improved use of a public good selects for the evolution of undifferentiated multicellularity. eLife 2:e00367. https://doi.org/10.7554/eLife.00367

Lande R, Arnold SJ (1983) The measurement of selection on correlated characters. Evolution 37:1210–1226. https://doi.org/10.2307/2408842

Lewontin RC (1970) The units of selection. Annu Rev Ecol Evol S 1:1–18

Libby E, Rainey PB (2013a) A conceptual framework for the evolutionary origins of multicellularity. Phys Biol 10:035001

Libby E, Rainey PB (2013b) Eco-evolutionary feedback and the tuning of proto-developmental life cycles. PLoS ONE 8:e82274

Lind PA, Libby E, Herzog J, Rainey PB (2019) Predicting mutational routes to new adaptive phenotypes. eLife 8:e38822. https://doi.org/10.7554/eLife.38822

Lloyd EA (1989) A structural approach to defining units of selection. Philos Sci 56:395–418. https://doi.org/10.2307/187992

Lloyd EA, Gould SJ (1993) Species selection on variability. Proc Natl Acad Sci USA 90:595–599. https://doi.org/10.1073/pnas.90.2.595

Maynard Smith J, Szathmáry E (1995) The Major Transitions in Evolution. Oxford University Press, Oxford

Michod RE (1999) Darwinian Dynamics, Evolutionary Transitions in Fitness and Individuality. Princeton University Press, Princeton, N.J.

Mikhailov KV, Konstantinova AV, Nikitin MA, et al (2009) The origin of Metazoa: a transition from temporal to spatial cell differentiation. Bioessays 31:758–768. https://doi.org/10.1002/bies.200800214

Okasha S (2006) Evolution and the Levels of Selection. Oxford University Press, Oxford

Okasha S (2012) Emergence, hierarchy and top-down causation in evolutionary biology. Interface Focus 2:49–54. https://doi.org/10.1098/rsfs.2011.0046

Rainey PB (2007) Unity from conflict. Nature 446:616–616

Rainey PB, de Monte S (2014) Resolving Conflicts During the Evolutionary Transition to Multicellular Life. Annu Rev Eco Evo Syst 45:599–620

Rainey PB, Kerr B (2010) Cheats as first propagules: A new hypothesis for the evolution of individuality during the transition from single cells to multicellularity. Bioessays 32:872–880. https://doi.org/Doi10.1002/Bies.201000039

Rainey PB, Rainey K (2003) Evolution of cooperation and conflict in experimental bacterial populations. Nature 425:72–74

Ratcliff WC, Denison RF, Borrello M, Travisano M (2012) Experimental evolution of multicellularity. Proc Natl Acad Sci USA 109:1595–1600. https://doi.org/10.1073/pnas.1115323109

Ratcliff WC, Herron MD, Howell K, et al (2013) Experimental evolution of an alternating uni-and multicellular life cycle in *Chlamydomonas reinhardtii*. Nat Commun 4:1–7. https://doi.org/10.1038/ncomms3742

Rose CJ (2020) Germ Lines and Extended Selection during the Evolutionary Transition to Multicellularity. In preparation

Rose CJ, Hammerschmidt K, Rainey PB (2020) Meta-population structure and the evolutionary transition to multicellularity. bioRxiv 407163. https://doi.org/10.1101/407163

Shelton DE, Michod RE (2014) Group selection and group adaptation during a major evolutionary transition: insights from the evolution of multicellularity in the volvocine algae. Biol Theory 1–18

Sober E, Lewontin RC (1982) Artifact, cause and genic selection. Philos Sci 49:157–180

Spiers AJ, Kahn SG, Bohannon J, et al (2002) Adaptive divergence in experimental populations of *Pseudomonas fluorescens*. I. Genetic and phenotypic bases of wrinkly spreader fitness. Genetics 161:33–46

Vrba E (1989) Levels of selection and sorting with special reference to the species level. Oxford surveys in evolutionary biology 6:111–168

Vrba E, Eldredge N (1984) Individuals, hierarchies and processes; towards a more complete evolutionary theory. Paleobiology 10:146–171

Wimsatt WC (1980) The units of selection and the structure of the multi-level genome. PSA: Proceedings of the Biennial Meeting of the Philosophy of Science Association 1980:122–183. https://doi.org/10.2307/192589

